# Genomic Mutations and Changes in Protein Secondary Structure and Solvent Accessibility of SARS-CoV-2 (COVID-19 Virus)

**DOI:** 10.1101/2020.07.10.171769

**Authors:** Thanh Thi Nguyen, Pubudu N. Pathirana, Thin Nguyen, Quoc Viet Hung Nguyen, Asim Bhatti, Dinh C. Nguyen, Dung Tien Nguyen, Ngoc Duy Nguyen, Douglas Creighton, Mohamed Abdelrazek

## Abstract

Severe acute respiratory syndrome coronavirus 2 (SARS-CoV-2) is a highly pathogenic virus that has caused the global COVID-19 pandemic. Tracing the evolution and transmission of the virus is crucial to respond to and control the pandemic through appropriate intervention strategies. This paper reports and analyses genomic mutations in the coding regions of SARS-CoV-2 and their probable protein secondary structure and solvent accessibility changes, which are predicted using deep learning models. Prediction results suggest that mutation D614G in the virus spike protein, which has attracted much attention from researchers, is unlikely to make changes in protein secondary structure and relative solvent accessibility. Based on 6,324 viral genome sequences, we create a spreadsheet dataset of point mutations that can facilitate the investigation of SARS-CoV-2 in many perspectives, especially in tracing the evolution and worldwide spread of the virus. Our analysis results also show that coding genes E, M, ORF6, ORF7a, ORF7b and ORF10 are most stable, potentially suitable to be targeted for vaccine and drug development.

## INTRODUCTION

Biological investigations of the novel coronavirus SARS-CoV-2 are important to understand the virus and help to propose appropriate responses to the pandemic. Scientists have been able to obtain genomic sequences of SARS-CoV-2 and have started analysis of these data. Reference genome of SARS-CoV-2 deposited to the National Center for Biotechnology Information (NCBI) GenBank sequence database (isolate Wuhan-Hu-1, accession number NC_045512) shows that SARS-CoV-2 is an RNA virus having a length of 29,903 nucleotides. Comparative genomic analysis results obtained in^1,2,3^ suggest that the COVID-19 virus may be originated in bats. Other studies show that pangolins may have served as the hosts for the virus^4,5^. Andersen et al.^6^ furthermore believe that SARS-CoV-2 is not a purposefully manipulated virus or constructed in a laboratory but has a natural origin. A study in^7^ using machine learning unsupervised clustering methods corroborates previous findings that SARS-CoV-2 belongs to the *Sarbecovirus* subgenus of the *Betacoronavirus* genus within the *Coronaviridae* family^8,9^. The whole genome analysis results also indicate that bats are more likely the reservoir hosts for the virus than pangolins. Another study in^10^ demonstrates that SARS-CoV-2 may have resulted from a recombination of a pangolin coronavirus and a bat coronavirus, and pangolins may have acted as an intermediate host for the virus.

Since the first cases were detected, the COVID-19 virus has spread to almost every country in the world and has been linked to the deaths of more than 404,000 people of over 7 million confirmed cases^11^. Tracing the evolution and spread of the virus is important for developing vaccines and drugs as well as proposing appropriate intervention strategies. Monitoring and analysing the viral genome mutations can be helpful for this task. Due to a strong immunologic pressure in humans, the virus may have mutated over time to circumvent responses of the human immune system. This leads to the creation of virus variants with possible different virulence, infectivity, and transmissibility^12^. This paper reports all point mutations occurring so far in SARS-CoV-2 and presents exemplified implications obtained from the analysis of these mutation pattern data. Four types of mutations, which include synonymous, nonsynonymous, insertion and deletion, are detected. We use 6,324 SARS-CoV-2 genome sequences collected in 45 countries and deposited to the NCBI GenBank so far and create a spreadsheet dataset of all mutations occurred across different genes. Eleven protein coding genes of SARS-CoV-2 have been identified, namely ORF1ab, spike (S), ORF3a, envelope (E), membrane (M), ORF6, ORF7a, ORF7b, ORF8, nucleocapsid (N) and ORF10. The order of these genes and their corresponding length are illustrated in Fig. 1.

**Fig 1.**
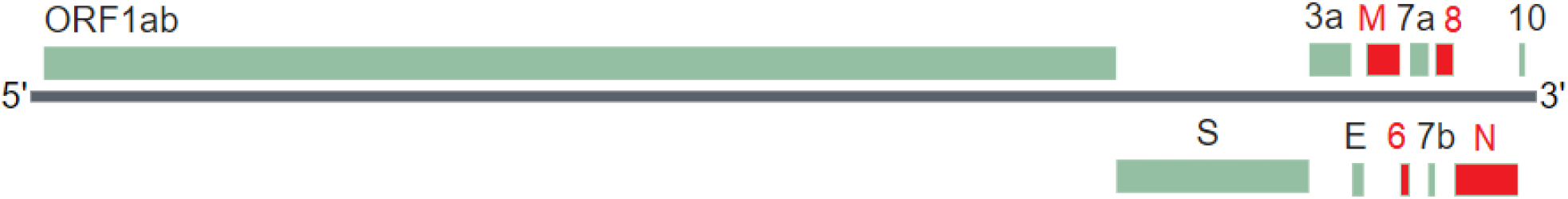
Protein coding genes of SARS-CoV-2, which consist of 7 nonstructural genes (ORF1ab, ORF3a, ORF6, ORF7a, ORF7b, ORF8 and ORF10) and 4 structural genes (S, E, M, and N). ORF1ab polyprotein is coded by gene ORF1ab at locations 266-21555 (based on the reference genome sequence NC_045512), surface glycoprotein coded by gene S at locations 21563-25384, ORF3a protein by gene ORF3a (25393-26220), envelope protein by gene E (26245-26472), membrane glycoprotein by gene M (26523-27191), ORF6 protein by gene ORF6 (27202-27387), ORF7a protein by gene ORF7a (27394-27759), ORF7b protein by gene ORF7b (27756-27887), ORF8 protein by gene ORF8 (27894-28259), nucleocapsid phosphoprotein by gene N (28274-29533), and ORF10 protein by gene ORF10 (29558-29674).

The genes S, E, M, and N produce structural proteins that play important roles in the virus functions. For example, the receptor-binding domain (RBD) region of the S protein can bind to a receptor of a host cell, e.g. the human and bat angiotensin-converting enzyme 2 (ACE2) receptor, enabling the entrance of the virus into the cell^13^. Predictions of protein structures may help understand the virus’ s functions and thus contribute to developing vaccines and therapeutics against the virus. In this paper, to evaluate the possible impacts of genomic mutations on the virus functions, we propose the use of the SSpro/ACCpro 5 methods to predict protein secondary structure and relative solvent accessibility^14^. These predictors were built using deep learning one-dimensional bidirectional recurrent neural networks incorporated in the SCRATCH-1D software suite (version 1.2, 2018)^15^. By comparing the prediction results obtained on the reference genome and mutated genomes, we are able to assess whether the detected mutations have the potential to change the protein structure and solvent accessibility, and thus lead to possible changes of the virus characteristics. Because of the functional importance of structural proteins, we only report the prediction results of these proteins in this study. The next section reviews related works in the literature. We then present materials and methods for SARS-CoV-2 mutation detection, and protein secondary structure and solvent accessibility prediction. Next we summarize statistics of SARS-CoV-2 mutations so far and implications of these mutations. Details of mutations in nonstructural ORF genes and structural S, E, M and N genes are presented after that. A full SARS-CoV-2 mutation spreadsheet report is provided in the Data Availability section.

## RELATED WORKS

Since the first genomes were collected in December 2019, there have been many findings on the mutations of SARS-CoV-2. For example, Phan^16^ analysed 86 genomes of SARS-CoV-2 downloaded from the the Global Initiative on Sharing All Influenza Data (GISAID) database (https://www.gisaid.org/) and found 93 mutations over the entire viral genome sequences. Among them, there are three mutations occurring in the RBD region of the spike surface glycoprotein S, including N354D, D364Y and V367F, with the numbers showing amino acid (AA) positions in the protein. That study also reveals three deletions in the genomes of SARS-CoV-2 obtained from Japan, USA and Australia. Two of these deletion mutations are in the ORF1ab polyprotein and one deletion occurs in the 3’ end of the genome. Likewise, a study in^17^ shows that the SARS-CoV-2 genomes may have undergone recurrent, independent mutations at 198 sites with 80% of these are of the nonsynonymous type. Alternatively, a SNP genotyping study in^18^ discovered highly frequent mutations in the genes encoding the S protein, RNA polymerase, RNA primase, and nucleoprotein. Those high-frequency SNP mutations are worth further investigations for vaccine development because they may be linked to the virus transmissibility and virulence. Tang et al.^19^ investigated 103 genomes of COVID-19 patients and discover mutations in 149 sites of these genomes. The study also shows that the spike gene S consistently has larger dS values (synonymous substitutions per synonymous site) than other genes. In addition, two major lineages of the virus, denoted as L and S, have been specified based on two tightly linked SNPs. The L lineage is found more prevalent than the S lineage among the examined sequences.

Korber et al.^20^ tracked the mutations of spike protein S of SARS-CoV-2 because it plays an important role in mediating infection of targeted cells and is the focus of vaccine and antibody therapy development efforts^21^. They detected 14 mutations in the spike protein that are growing, especially the mutation D614G that rapidly becomes the dominant form when spread to a new geographical region. Likewise, Hashimi^22^ analysed the mutation frequency in the spike protein S of 796 SARS-CoV-2 genomes downloaded from the GISAID and GenBank databases. The study found 64 mutations occurring in the S protein sequences obtained from multiple countries. It suggests that the virus is spreading in two forms, the D614 form (residue D at position 614 in the S protein) takes 68.5% while the G614 form takes 31.5% proportion of the examined isolates. Koyama et al.^23^ on the other hand found several variants of SARS-CoV-2 that may cause drifts and escape from immune recognition by using the prediction results of B-cell and T-cell epitopes in^24^. Typically, the mutation D614G occurring in the spike protein is found prevalent in the European population. This mutation may have caused antigenic drift, resulting in vaccine mismatches that lead to a high mortality rate of this population.

A recent situation report^25^ by Nextstrain^26^ on genomic epidemiology of novel coronavirus using 5,193 publicly shared COVID-19 genomes shows that SARS-CoV-2 on average accumulates changes at a rate of 24 substitutions per year. This is approximately equivalent to 1 mutation per 1,000 bases in a year. This evolutionary rate of SARS-CoV-2 is typical for a coronavirus, and it is smaller than that of influenza (average 2 mutations per 1,000 bases per year) and HIV (average 4 mutations per 1,000 bases per year). Shen et al.^12^ conducted metatranscriptome sequencing for bronchoalveolar lavage fluid samples obtained from 8 patients with COVID-19 and found no evidence for the transmission of intrahost variants as well as a high evolution rate of the virus with the number of intrahost variants ranged from 0 to 51 around a median number of 4. Pachetti et al.^27^ examined 220 genomic sequences of COVID-19 patients from the GISAID database and discovered 8 novel recurrent mutations at nucleotide locations 1397, 2891, 14408, 17746, 17857, 18060, 23403 and 28881. Mutations at locations 2891, 3036, 14408, 23403 and 28881 are mostly found in Europe while those at locations 17746, 17857 and 18060 occur in sequences obtained from patients in North America. Likewise, a study in^28^ on 95 SARS-CoV-2 complete genome sequences discovered 116 mutations. Among them, the mutations at position C8782T in the ORF1ab gene, T28144C in the ORF8 gene and C29095T in the N gene are common.

## MATERIALS AND METHODS

### SARS-CoV-2 Mutation Detection

We use 6,324 sequence records downloaded from the NCBI GenBank database on 2020-06-17. The latest collection date for the samples from which the sequences were derived was on *2020-06-05*. The data, which were collected in 45 countries, include both nucleotide sequences and protein translations of coding genes. A proportion of the 6,324 records have sequences of only few proteins, i.e. these records do not annotate all 11 proteins (ORF1ab, ORF3a, ORF6, ORF7a, ORF7b, ORF8, ORF10, S, E, M and N). The number of available sequences is thus different from one protein to another (see column “Avai Num” in Table 1). Genome sequences that do not specify country or AA sequences that contain letter “X” representing an unknown AA are excluded in our calculations. We use the genome obtained from the isolate *Wuhan-Hu-1, accession number NC_045512* as the reference genome. For the mutation detection purpose, we apply a dynamic programming algorithm to protein AA sequences to get global pairwise alignments between a reference sequence and a query sequence. Specifically, we use the Python Bio.pairwise2.align.globalms function (https://biopython.org/docs/dev/api/Bio.pairwise2.html) where a match is given 2 points, a mismatch is deducted 0.5 points, 2 points are deducted when opening a gap, and 1 point is deducted when extending it. Gaps are then inserted into nucleotide sequences corresponding to the resulted protein sequence alignments. Using the resulted pairwise alignments, we are able to compare query sequences and the reference sequences at each position and identify locations of insertion, deletion, synonymous and nonsynonymous mutations.

**Table 1.** Summary of SARS-CoV-2 mutations on each protein and secondary structure and relative solvent accessibility changes

| Proteins | AA Length | Avai. Num | No. Mu | No. Mu Rate | Delete | Insert | Nonsyn | Syn | Nonsyn/Syn | Struct Change | Acc Change |
| --- | --- | --- | --- | --- | --- | --- | --- | --- | --- | --- | --- |
| ORF1ab | 7096 | 3726 | 68 | 0.02 | 170 | 6 | 8330 | 6540 | 1.27 | NA | NA |
| S | 1273 | 4434 | 872 | 0.20 | 34 | 0 | 3711 | 670 | 5.54 | 164 | 184 |
| ORF3a | 275 | 5527 | 2182 | 0.39 | 6 | 1 | 3400 | 277 | 12.27 | NA | NA |
| E | 75 | 5852 | 5747 | 0.98 | 0 | 0 | 28 | 77 | 0.36 | 18 | 7 |
| M | 222 | 5677 | 5128 | 0.90 | 0 | 0 | 121 | 444 | 0.27 | 30 | 8 |
| ORF6 | 61 | 5792 | 5657 | 0.98 | 18 | 2 | 72 | 64 | 1.12 | NA | NA |
| ORF7a | 121 | 5321 | 5164 | 0.97 | 15 | 0 | 106 | 56 | 1.89 | NA | NA |
| ORF7b | 43 | 5175 | 5120 | 0.99 | 0 | 0 | 24 | 36 | 0.67 | NA | NA |
| ORF8 | 121 | 5732 | 4292 | 0.75 | 0 | 8 | 1563 | 74 | 21.12 | NA | NA |
| N | 419 | 5281 | 4051 | 0.77 | 15 | 0 | 1927 | 392 | 4.92 | 1678 | 37 |
| ORF10 | 38 | 5891 | 5829 | 0.99 | 0 | 0 | 19 | 43 | 0.44 | NA | NA |

### Secondary Structure and Solvent Accessibility Prediction

Virus protein structure plays a key role in its functions and a change in structure shape may affect its functions, virulence, infectivity and transmissibility, possibly resulting in non-functional proteins. Protein secondary structure is defined by hydrogen bonding patterns, which make an intermediate form before the protein folds into a three-dimensional shape composing its tertiary structure. Eight types of protein secondary structure defined by the Dictionary of Protein Secondary Structure (DSSP) include 3^10^ helix (G), *α* helix (H), *π* helix (I), hydrogen bonded turn (T), extended strand in parallel and/or anti-parallel *β*-sheet conformation (E), residue in isolated *β*-bridge (B), bend (S) and coil (C). The DSSP tool assigns every residue to one of the eight possible states. In a reduced form, these 8 conformational states can be diminished to 3 states: H = {H, G, I}, E = {E, B} and C = {S, T, C}^29^. The protein secondary structure represents interactions between neighboring or near-by AAs as its functional three-dimensional shape is created through the polypeptide folding. We thus determine a change in protein secondary structure if any change happens in the structures of the mutated AA and its 10 neighboring AAs compared to those of the reference sequence. In detail, we consider 5 AAs ahead and 5 AAs behind the mutated AA. The same approach is applied when considering a change of the protein relative solvent accessibility. Solvent-exposed area represents the area of a biomolecule on a surface that is accessible to a solvent. Accordingly, a residue is considered as exposed if at least 25% of that residue must be exposed, denoted as the “e” state. Alternatively, the residue is determined as buried, i.e. the “b” state. There have been various protein secondary structure prediction programs in the literature and many of those were developed based on artificial intelligence models using protein AA sequences such as JPred4^30^, Spider2^31^, Porter 5^32^, RaptorX^33^, PSSpred^34^, YASSPP^35^ and SSpro^14^. In this paper, we use the protein secondary structure and relative solvent accessibility prediction methods SSpro/ACCpro 5^14^ within the SCRATCH-1D software suite (release 1.2, 2018)^15^. These predictors were built using the bidirectional recursive neural networks and a combination of the sequence similarity and sequence-based structural similarity to sequences in the Protein Data Bank^36^. Prediction results of 8-class structure (SSpro8 predictor) and 25%-threshold relative solvent accessibility (ACCpro predictor) are used for statistics on protein secondary structure and accessibility changes. We however also report in the spreadsheet supplemental information prediction results of 3-class structure (SSpro predictor) and relative solvent accessibility on 20 thresholds, ranging from 0% to 95% with a 5% step (the ACCpro20 predictor within the SCRATCH-1D software).

## SUMMARY OF SARS-COV-2 MUTATIONS

Table 1 summarizes statistics of SARS-CoV-2 mutations so far. “AA Length” indicates the length of the protein AA sequence derived from the SARS-CoV-2 reference genome. “Avai. Num” denotes the number of records among 6,324 NCBI GenBank records that have the complete sequence of the corresponding protein. “No. Mu” refers to the number of sequences that do not have any mutations compared to the reference sequence. “No. Mu Rate” is the ratio between “No. Mu” and “Avai. Num”. “Delete” means the number of deletion mutations occurring in the AA sequences of the protein. This number may be larger than the number of sequences having deletion mutations because an AA sequence may have more than one deletion. Likewise, “Insert”, “Nonsyn” and “Syn” show the number of insertion, nonsynonymous and synonymous mutations occurring in the protein AA sequences. “Nonsyn/Syn” demonstrates a ratio between the number of nonsynonymous mutations versus the number of synonymous mutations. “Struct Change” means the number of *nonsynonymous mutations* that have protein secondary structure change potential based on the SSpro8 predictor of the SCRATCH-1D software. Similarly, “Acc Change” refers to the number of nonsynonymous mutations that have potential to change the protein relative solvent accessibility based on the ACCpro predictor of the SCRATCH-1D software. Insertion and deletion mutations alter protein secondary structure and solvent accessibility by default so that they are not included in the structure and solvent accessibility change statistics.

Table 1 shows that the ORF3a and ORF8 proteins have the number of nonsynonymous mutations significantly larger than that of the synonymous mutations. In contrast, this ratio in proteins E, M, ORF7b and ORF10 are very small (less than 1). These proteins could be targeted for vaccine and drug development as they have less variations than other proteins. These findings are supported by results presented in Figs. 2-4 where we plot the number of insertion, deletion and nonsynonymous mutations against different locations in the proteins. A spike in these figures demonstrates a large number of insertion, deletion and nonsynonymous mutations. Regions between spikes are stable and can be useful for further research for vaccine and drug development. For example, in protein ORF1ab (Fig. 2), regions [1…264], [266…3605], [3607…4714], [4716…5827], [5829…5864] and [5866…7096] are relatively stable. In protein S (Fig. 3), entire regions before and after the spike at position 614 are almost unchanged. Fig. 4 presents variations of multiple proteins. In addition to proteins E, M, ORF7b and ORF10, we find from Fig. 4 that proteins ORF6 and ORF7a are also relatively stable without a large number of variations at any particular locations. This is justified by data in the column “No. Mu Rate” in Table 1, which shows the ratio between “No. Mu” and “Avai. Num”, i.e. the ratio between the number of sequences having no mutations in a specific protein versus the total number of available sequences of that protein. Proteins E, M, ORF6, ORF7a, ORF7b and ORF10 have a large number of sequences with no mutations, therefore their ratios are very large, respectively of 0.98, 0.90, 0.98, 0.97, 0.99, and 0.99. This is because the size of these proteins is relatively small (see data in the column “AA Length” in Table 1). These proteins are considered as most stable when compared to other proteins, which have the ratios of less than 0.8. These results are consistent with the data shown in Fig. 2-4.

**Fig 2.**
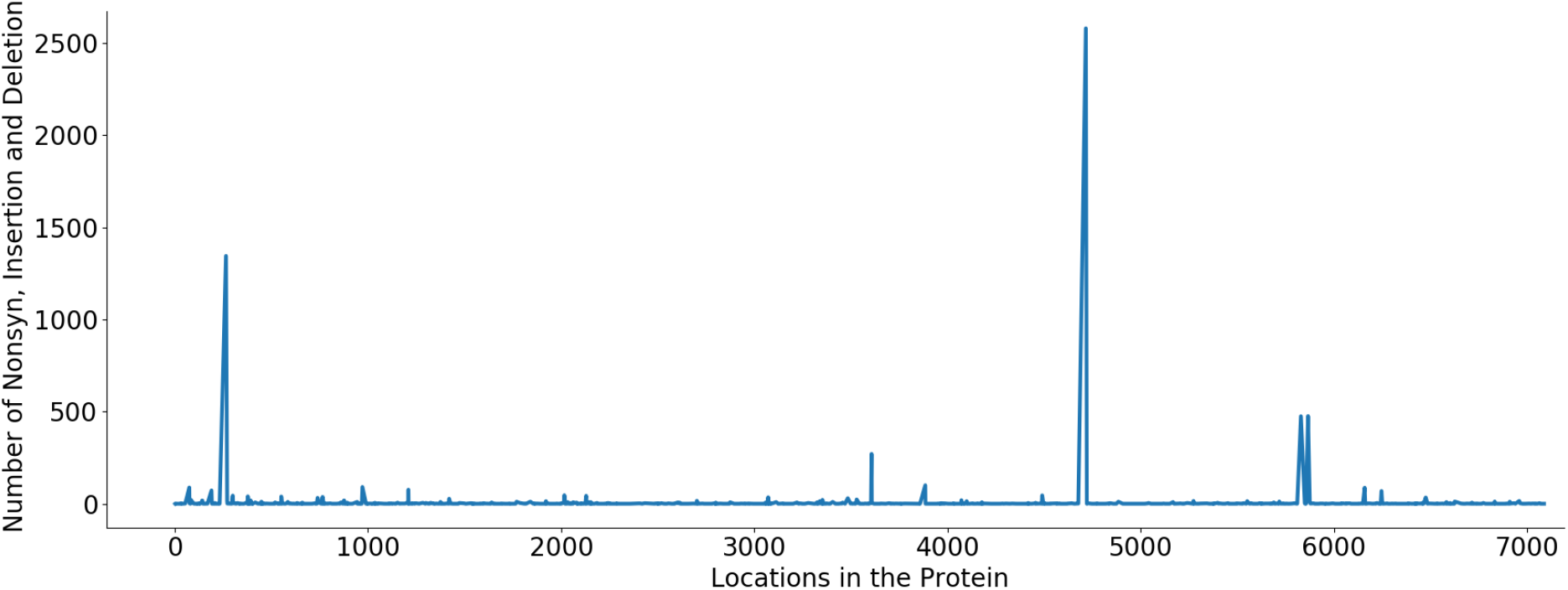
Protein ORF1ab - The number of insertion, deletion and nonsynonymous mutations at different locations in the protein. Spikes at locations: T265I (1344), L3606F (271), P4715L (2576), P5828L (475) and Y5865C (476). Regions between these spikes are stable and could be targeted for vaccine and drug development.

**Fig 3.**
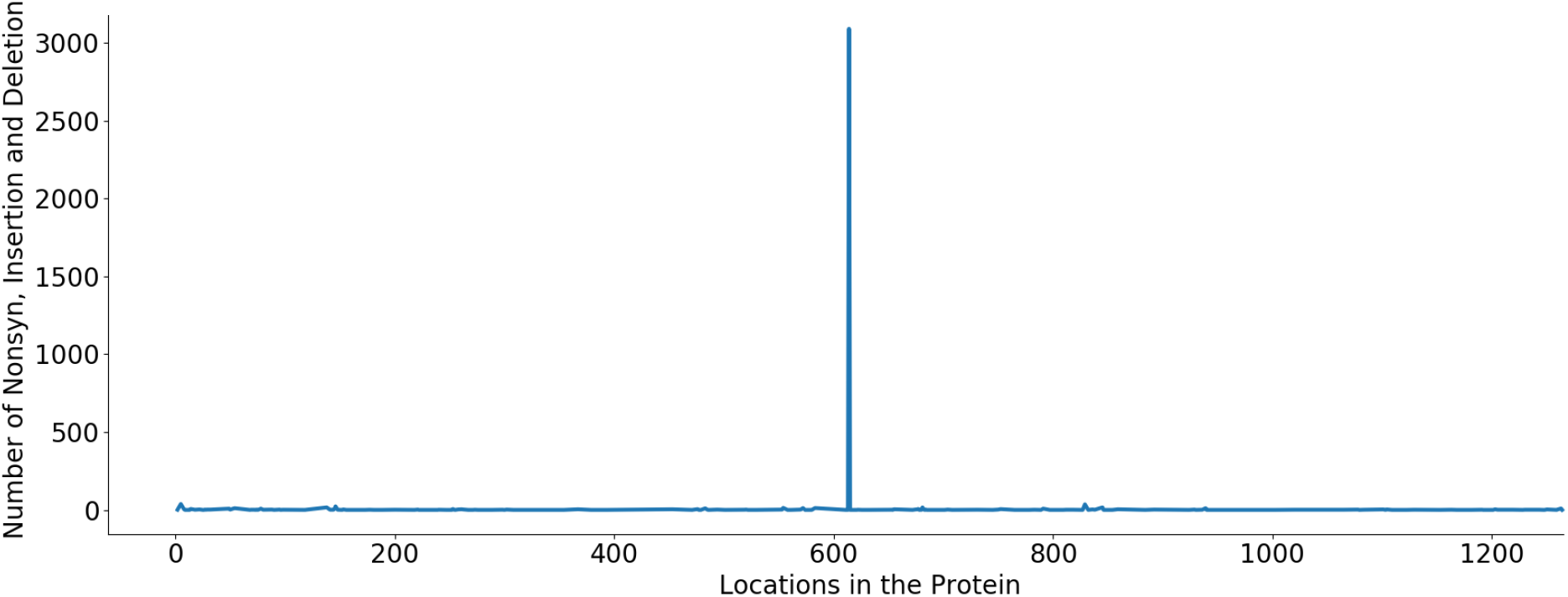
Protein S - The number of insertion, deletion and nonsynonymous mutations at different locations in the protein. A spike at location D614G (3089) while other regions of the protein are stable.

**Fig 4.**
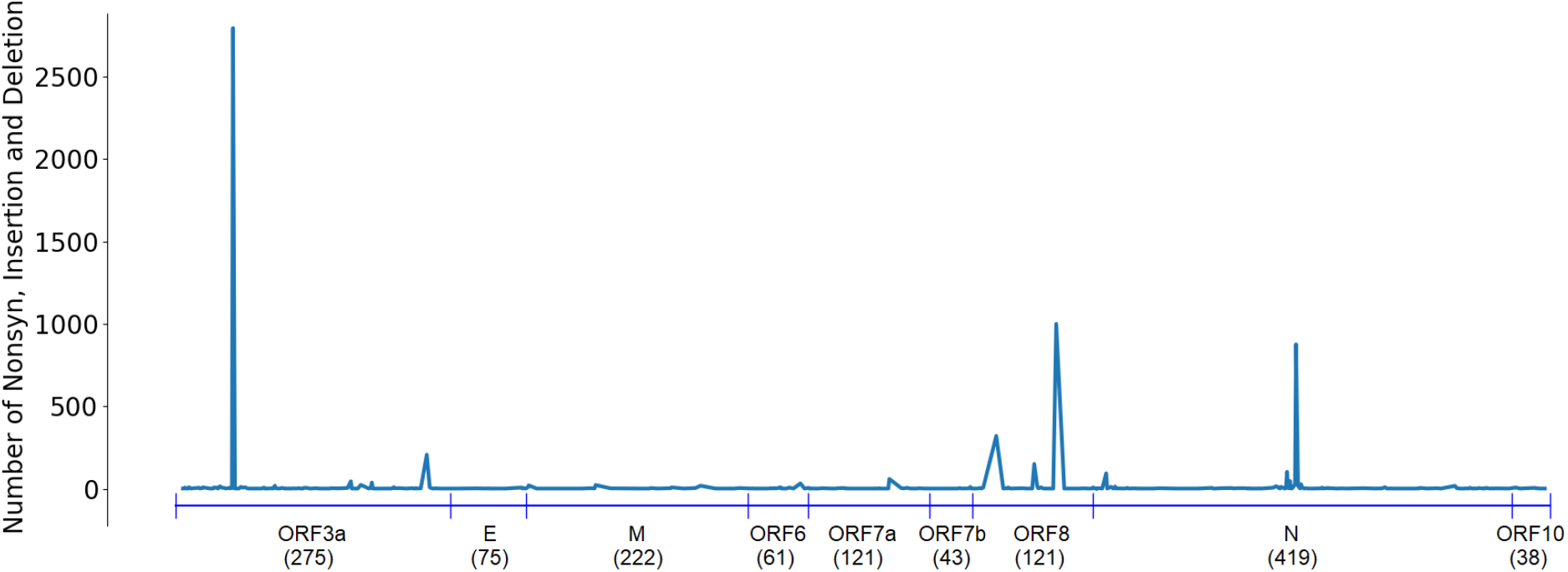
The number of insertion, deletion and nonsynonymous mutations at different locations in the proteins ORF3a, E, M, ORF6, ORF7a, ORF7b, ORF8, N and ORF10. The number below protein names are the length of that protein. Protein ORF3a: two spikes at Q57H (2795) and G251V (206); protein ORF8: two spikes at S24L (320) and L84S (1000); protein N: R203K (876) and G204R (433); Other proteins E, M, ORF6, ORF7a, ORF7b and ORF10 are almost entirely stable.

Protein N has 1,927 nonsynonymous mutations but 1,678 of them are likely to make changes in protein secondary structure, making a ratio of 87.08%. This is considerably larger than those of protein S (4.42%), protein M (24.79%) and protein E (64.29%). The number of solvent accessibility changes of protein S is larger than its structure changes: 184 vs 164. This however is opposite in other structural proteins: E (7 vs 18), M (8 vs 30) and N (37 vs 1,678).

## DETAILS OF MUTATIONS IN NONSTRUCTURAL ORF GENES

### Gene ORF1ab

The ORF1ab polyprotein has 7,096 AAs. Among 6,324 records deposited to the NCBI GenBank database, only 3,726 genomes have the complete coding sequence (CDS) of protein ORF1ab, with 1,024 unique AA sequences. This is quite a large number compared to other proteins but understandable because ORF1ab is the longest protein of SARS-CoV-2 and thus has a large number of variations. Only 119 sequences have no mutation or synonymous mutations while the rest 3,607 sequences have insertion, deletion or nonsynonymous mutations. There are *two distinct insertion mutations* −3603F and −7041F occurring in six different sequences. The insertion −3603F occurs in five sequences: MT507793 (collected in Jamaica on 2020-03-11), MT614545 (USA: NY on 2020-03-16), MT451423 (Australia: Victoria on 2020-03-28), MT451433 (Australia: Victoria on 2020-03-29) and MT451522 (Australia: Victoria on 2020-03-30). The insertion −7041F occurs only in MT188341, collected in USA: MN on 2020-03-05. There are total 170 deletion mutations, with *48 distinct deletions*. Deletions occurring in 10 or more sequences are reported in Table 2. Fig. 5 shows an alignment of sequences having a large number of deletions in the ORF1ab protein. Alternatively, 8,330 nonsynonymous mutations are found, in which 1,067 mutations are distinct. Table 3 presents common nonsynonymous mutations (those occurring in 50 or more sequences) in the ORF1ab protein. Details of other mutations are reported in the spreadsheet dataset in the Data Availability section. Notable mutations are P4715L occurring in 2,576 sequences and T265I occurring in 1,344 sequences.

**Table 2.** Gene ORF1ab − Deletion mutations occurring in 10 or more sequences

| Mu | Total | First Date | First Country | Latest Date | Latest Country | Distribution |
| --- | --- | --- | --- | --- | --- | --- |
| M85- | 21 | 2020-01-27 | USA: CA | 2020-04-27 | USA | USA (17) Greece (2) Australia (2) |
| V86- | 16 | 2020-01-27 | USA: CA | 2020-04-07 | USA | USA (12) Greece (2) Australia (2) |
| K141- | 18 | 2020-01-27 | USA: CA | 2020-04-08 | USA: CA | USA (18) |
| S142- | 18 | 2020-01-27 | USA: CA | 2020-04-08 | USA: CA | USA (18) |
| F143- | 18 | 2020-01-27 | USA: CA | 2020-04-08 | USA: CA | USA (18) |
| D448- | 12 | 2020-03-07 | USA: NV | 2020-04-29 | Netherlands | Netherlands (6) USA (4) Pakistan (1) Australia (1) |

**Table 3.** Gene ORF1ab - Nonsynonymous mutations occurring in 50 or more sequences

| Mu | Total | First Date | First Country | Latest Date | Latest Country | Distribution |
| --- | --- | --- | --- | --- | --- | --- |
| D75E | 89 | 2020-01-28 | USA: IL | 2020-05-11 | USA | USA (68) Australia (20) Taiwan (1) |
| F190L | 73 | 2020-03-12 | USA: WA | 2020-04-14 | USA | USA (72) Australia (1) |
| T265I | 1344 | 2020-02-26 | Taiwan | 2020-05-26 | Australia: Victoria | USA (1152) Australia (136) France (18) Germany (16) Taiwan (9) Czech Republic (4) Puerto Rico (3) Jamaica (2) Tunisia (1) India (1) Greece (1) Colombia (1) |
| P971L | 90 | 2020-01-28 | USA: IL | 2020-05-11 | USA | USA (68) Australia (20) Taiwan (2) |
| E1209D | 75 | 2020-04-01 | USA | 2020-04-06 | USA | USA (75) |
| L3606F | 271 | 2020-01-17 | China: Yunnan | 2020-05-27 | Australia: Victoria | USA (107) Australia (69) India (15) Greece (12) Taiwan (11) France (11) China (6) Hong Kong (5) Czech Republic (4) Timor-Leste (3) South Korea (3) Japan (3) Jamaica (3) Guam (3) Malaysia (2) Brazil (2) Turkey (1) Tunisia (1) Thailand (1) Sri Lanka (1) Spain (1) Pakistan (1) Kenya (1) Kazakhstan (1) Italy (1) Israel (1) Iran (1) Bangladesh (1) |
| S3884L | 101 | 2020-03-05 | USA: NY | 2020-04-30 | USA: CA | USA (98) Australia (2) Puerto Rico (1) |
| P4715L | 2576 | 2020-02 | Germany | 2020-06-05 | India: Vadodara | USA (1684) Australia (362) India (184) Greece (62) France (59) Bangladesh (48) Japan (27) Germany (27) Czech Republic (21) Poland (18) Taiwan (13) Spain (10) Netherlands (8) Serbia (7) Egypt (7) Italy (6) Turkey (4) Tunisia (4) Puerto Rico (4) China (3) Thailand (2) Sri Lanka (2) Russia (2) Kazakhstan (2) Jamaica (2) Hong Kong (2) South Africa (1) Peru (1) Nigeria (1) Morocco (1) Israel (1) Colombia (1) |
| P5828L | 475 | 2020-02-20 | USA: WA | 2020-04-30 | USA: CA | USA (443) Australia (31) Puerto Rico (1) |
| Y5865C | 476 | 2020-02-20 | USA: WA | 2020-04-30 | USA: CA | USA (444) Australia (31) Puerto Rico (1) |
| F6158L | 88 | 2020-03 | USA: VA | 2020-05-11 | USA | USA (67) Australia (20) Taiwan (1) |
| A6245V | 70 | 2020-03-05 | USA: NY | 2020-04-30 | USA: CA | USA (68) Puerto Rico (1) Australia (1) |

**Fig 5.**
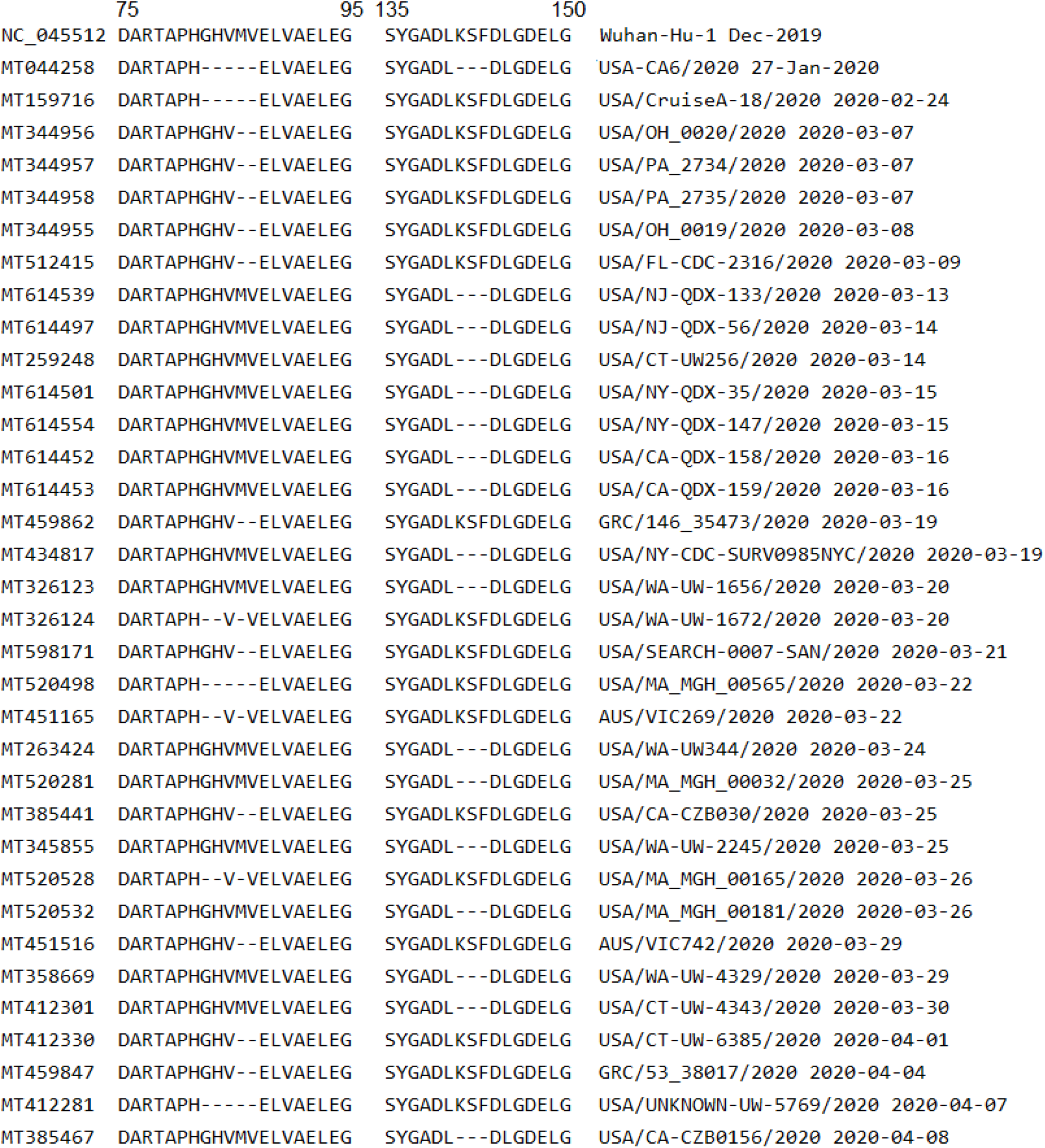
Alignment of sequences having deletion mutations at positions M85-, V86- or K141-, S142-, F143-, which are major deletions in the ORF1ab protein (Table 2). The GenBank accession numbers are presented on the left while isolate names and collected dates are on the right. The numbers on top show the positions of AAs in the protein and isolates are ordered by collected dates. The first isolate having these deletions is USA-CA6/2020 (record MT044258 in second row), collected on 2020-01-27 in USA: CA. This is also the isolate having the largest number of deletions: five sequentially at G82-, H83-, V84-, M85-, V86- and three at K141-, S142-, F143-. The other patients followed were possibly infected by this first case but more data such as travel history are needed to confirm this hypothesis.

### Gene ORF3a

The ORF3a protein has 275 AAs with its complete CDS appearing in 5,527 isolates (146 unique AA sequences). Among these, 2,321 sequences have no mutation or only synonymous mutations, and 3,206 sequences have insertion, deletion or nonsynonymous mutations. *One insertion mutation* occurs in sequence MT449656, collected in USA: CA on 2020-04-13, at position-230F. There are *six distinct deletions*, three of them occur sequentially in MT293186, collected in USA: WA on 2020-03-17: I10-, G11- and T12-. Three other deletions occur in three different sequences: MT326059 (collected in USA on 2020-03-24), MT358717 (USA: WA on 2020-03-27) and MT474130 (USA: CA on 2020-03-31) at positions V48-, V256- and N257-, respectively. A total of 3,400 nonsynonymous mutations (125 unique) are found in the ORF3a protein. Nonsynonymous mutations occur in 10 or more sequences are reported in Table 4. Notably, the mutation *Q57H occurs in 2,795 sequences* collected in many countries. This is an emerging and active mutation, which requires further investigation as the latest case of this mutation was on 2020-06-05, same as the latest collection date of the entire downloaded dataset. The mutation *G251V occurring in 206 sequences* is also a prevalent mutation in the ORF3a protein.

**Table 4.** Gene ORF3a - Nonsynonymous mutations occurring in 10 or more sequences

| Mu | Total | First Date | First Country | Latest Date | Latest Country | Distribution |
| --- | --- | --- | --- | --- | --- | --- |
| G44V | 14 | 2020-03-17 | USA: WA | 2020-04-09 | USA: AK | USA (14) |
| Q57H | 2795 | 2020-02-29 | USA: NH | 2020-06-05 | India: Gandhinagar | USA (2272) Australia (300) India (116) France (29) Germany (22) Egypt (10) Puerto Rico (8) Greece (8) Taiwan (6) Japan (5) Czech Republic (5) Tunisia (4) Bangladesh (4) Jamaica (3) Serbia (1) Kenya (1) Colombia (1) |
| L65F | 11 | 2020-04 | USA: VA | 2020-04 | USA: VA | USA (11) |
| T175I | 45 | 2020-03-09 | USA: MI | 2020-05-23 | Bangladesh | USA (39) Australia (3) India (1) Greece (1) Bangladesh (1) |
| Q185H | 23 | 2020-04 | USA: Virginia | 2020-04-30 | USA: CA | USA (23) |
| G196V | 36 | 2020-02-26 | Spain: Valencia | 2020-04-07 | Australia: Victoria | Australia (26) Spain (4) USA (3) Greece (2) Kazakhstan (1) |
| G251V | 206 | 2020-01-22 | USA: CA | 2020-04-22 | Australia: Victoria | Australia (84) USA (67) Greece (13) France (8) Thailand (5) Hong Kong (5) South Korea (4) Jamaica (4) China (3) Timor-Leste (2) Italy (2) Germany (2) Brazil (2) Taiwan (1) Sweden (1) Sri Lanka (1) Spain (1) Kenya (1) |

### Gene ORF6

The ORF6 protein has 61 AAs, appearing in 5,792 isolates with 25 unique AA sequences. Among these, 5,719 sequences have no mutation or only synonymous mutations and 73 sequences have insertion, deletion or nonsynonymous mutations. *Two insertion mutations* occur in record MT520188 at positions −62R and −63T (end of the sequence). *Nine continual deletions* occur similarly in 2 sequences: MT547814 (collected in Hong Kong on 2020-01-22 from an adult male patient^37^) and MT609561 (USA: Virginia in 2020-04). These deletions are F22-, K23-, V24-, S25-, I26-, W27-, N28-, L29- and D30-. Alignment of these sequences with the reference genome is displayed in Fig. 6. The isolate MT547814 thus may have transmitted the virus to MT609561 but this implication needs to be corroborated by patients’ travel history. There are 23 distinct nonsynonymous mutations and those occurring in 2 or more sequences are presented in Table 5.

**Table 5.**
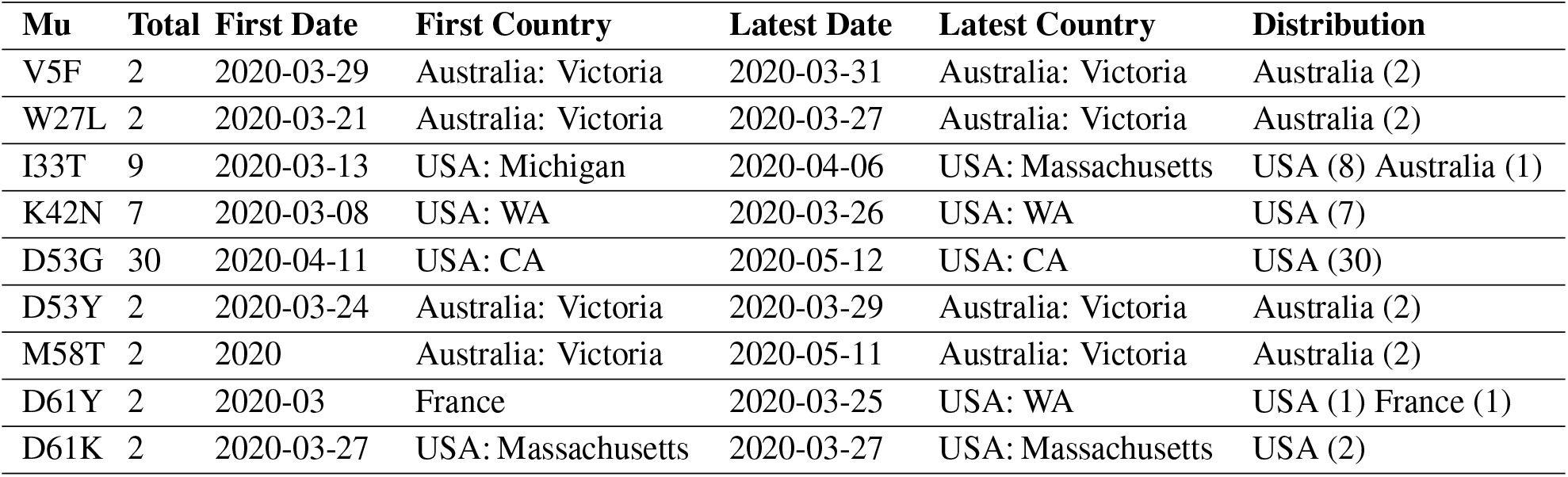
Gene ORF6 - Nonsynonymous mutations occurring in 2 or more sequences

**Fig 6.**
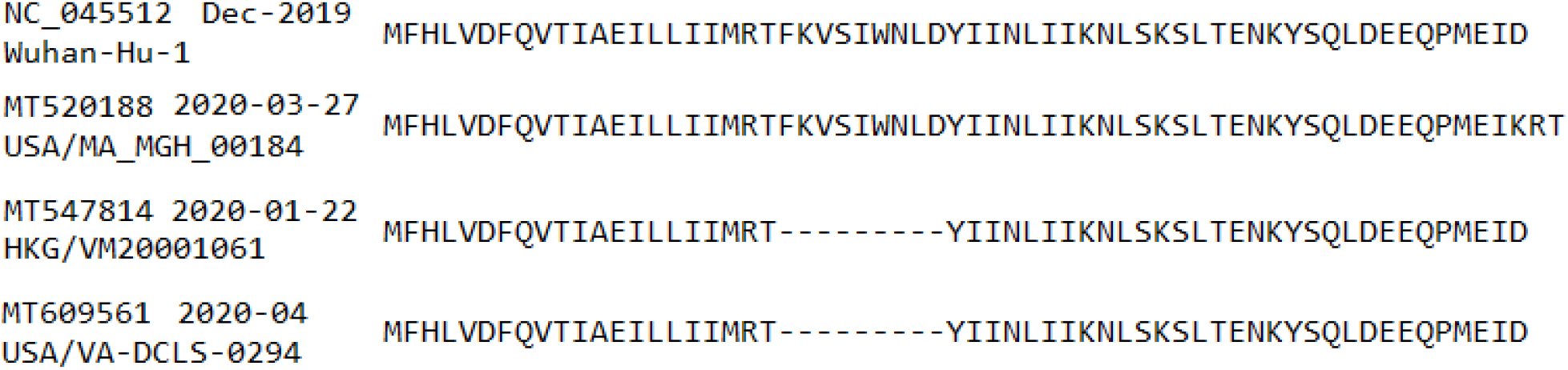
Insertion and deletion mutations in protein ORF6. The GenBank accession numbers, collected dates and isolate names are presented on the left. One synonymous mutation D61K and two insertions −62R and −63T at the end of isolate USA/MA_MGH_00184/2020 (MT520188) is interesting while there is a high chance that HKG/VM20001061/2020 has spread to USA/VA-DCLS-0294/2020.

### Gene ORF7a

The ORF7a protein has 121 AAs in length, found in 5,321 isolates with 34 unique AA sequences. Among these, 5,215 sequences have no mutations or only synonymous mutations, while the rest 106 sequences have deletion or nonsynonymous mutations. *No insertion mutation* is found in gene ORF7a. There are *15 deletion mutations* occurring in 2 records: MT520425 (collected in USA: Massachusetts on 2020-03-27) and MT507795 (USA on 2020-04-06). The MT520425 sequence has 1 deletion at position L77-while the MT507795 sequence has 14 sequential deletions F63-, A64-, F65-, A66-, C67-, P68-, D69-, G70-, V71-, K72-, H73-, V74-, Y75- and Q76-. Alignment of these sequences with that of the reference genome is shown in Fig. 7. There are 32 distinct nonsynonymous mutations with those occurring in 2 or more sequences are reported in Table 6.

**Table 6.** Gene ORF7a - Nonsynonymous mutations occurring in 2 or more sequences

| Mu | Total | First Date | First Country | Latest Date | Latest Country | Distribution |
| --- | --- | --- | --- | --- | --- | --- |
| T14I | 3 | 2020-03-17 | USA | 2020-05-14 | Australia: Victoria | Australia (2) USA (1) |
| V29L | 3 | 2020-03-25 | Australia: Victoria | 2020-04-06 | USA: Houma, LA | Australia (2) USA (1) |
| S36F | 4 | 2020-04-04 | USA: CA | 2020-04-26 | USA: CA | USA (4) |
| V71I | 2 | 2020-03-20 | USA: WA | 2020-03-28 | USA: WA | USA (2) |
| S81L | 59 | 2020-03-07 | Australia: Victoria | 2020-04-15 | USA: Illinois | Australia (35) USA (24) |
| V93F | 3 | 2020-03-24 | Australia: Victoria | 2020-04-01 | Australia: Victoria | Australia (3) |
| E95K | 2 | 2020-06-03 | India: Ahmedabad | 2020-06-03 | India: Ahmedabad | India (2) |
| P99S | 4 | 2020-03 | USA: VA | 2020-03-31 | Australia: Victoria | USA (3) Australia (1) |
| I110T | 3 | 2020-03-14 | USA: WA | 2020-03-20 | USA | USA (3) |

**Fig 7.**
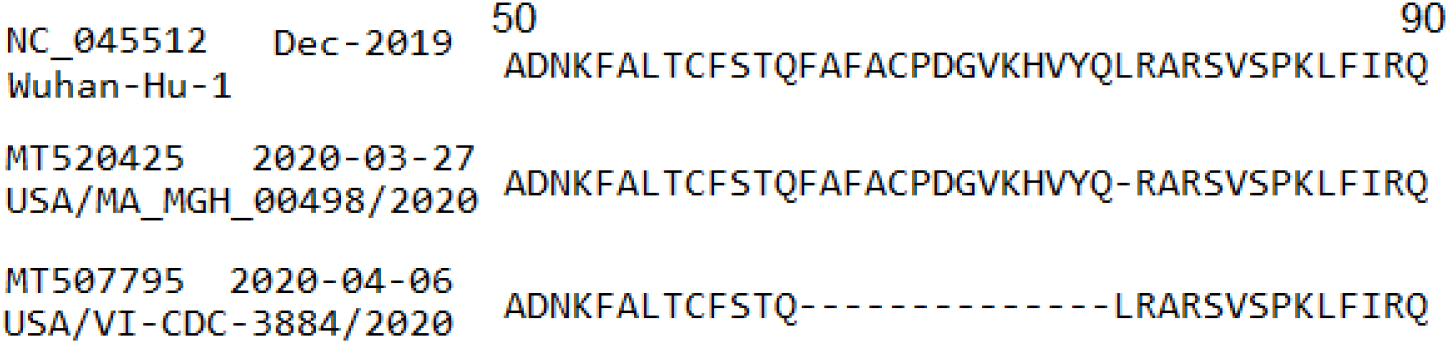
Deletions in protein ORF6 with the GenBank accession numbers, collected dates and isolate names presented on the left. The large 14 sequential deletions in the isolate USA/VI-CDC-3884/2020 (MT507795) are worth a further study as its patient’ s clinical data may show some difference with other COVID-19 patients.

### Gene ORF7b

The ORF7b protein has 43 AAs with its complete CDS appearing in 5,175 isolates, forming a set of 11 unique AA sequences. There are 5,151 sequences having no mutations or only synonymous mutations and 24 sequences having nonsynonymous mutations. *No insertion or deletion mutations* are found in gene ORF7b. This along with a small number of nonsynonymous mutations indicate that ORF7b is a stable gene. Distinct nonsynonymous mutations (10 of them) include F19L, F28Y, F30L, S31L, L32F, T40I, C41F, C41S, H42Y and A43T. Summary of nonsynonymous mutations in gene ORF7b occurring in 2 or more sequences is shown in Table 7.

**Table 7.** Gene ORF7b - Nonsynonymous mutations occurring in 2 or more sequences

| Mu | Total | First Date | First Country | Latest Date | Latest Country | Distribution |
| --- | --- | --- | --- | --- | --- | --- |
| F30L | 3 | 2020-04-26 | USA: CA | 2020-04-30 | USA: CA | USA (3) |
| S31L | 3 | 2020-03-22 | USA: Michigan | 2020-05-09 | India: Botad | USA (2) India (1) |
| T40I | 2 | 2020-03-24 | USA: CA | 2020-04-08 | USA: Michigan | USA (2) |
| C41F | 10 | 2020-03-01 | Thailand | 2020-04-14 | Australia: Victoria | Thailand (5) Australia (5) |

### Gene ORF8

This gene codes the ORF8 protein that has a length of 121 AAs. There are 5,732 isolates containing complete CDS of gene ORF8 with 55 unique AA sequences. Among 5,732 obtained sequences, 4,346 of them have no mutation or only synonymous mutations and the rest 1,386 have mutations either insertions or nonsynonymous mutations. *No deletion mutations* are found in gene ORF8. *Four distinct insertion mutations* occur similarly in 2 sequences: MT568638 (collected in China: Guangzhou on 2020-02-25) and MT507032 (USA: FL on 2020-04-21). These insertions are at the end of the protein: −122K, −123R, −124T and −125N. Alignment of sequences having insertions with the reference genome is shown in Fig. 8. The number of nonsynonymous mutations in gene ORF8 is 1,563 with 50 distinct ones. Summary of nonsynonymous mutations occurring in 2 or more sequences is presented in Table 8. Notable mutation in this gene is *L84S*, which occurs in 1,000 sequences with the first case collected *at the beginning of the pandemic* in China: Wuhan on 2019-12-30 and the latest case in Australia: Victoria on 2020-05-27.

**Table 8.** Gene ORF8 - Nonsynonymous mutations occurring in 2 or more sequences

| **Mu** | **Total** | **First Date** | **First Country** | **Latest Date** | **Latest Country** | **Distribution** |
| --- | --- | --- | --- | --- | --- | --- |
| G8R | 3 | 2020-04-05 | USA: FL | 2020-04-17 | USA | USA (3) |
| T11K | 2 | 2020-04-14 | USA: CA | 2020-05-12 | USA: CA | USA (2) |
| T11A | 4 | 2020-03-23 | USA | 2020-03-24 | USA: WA | USA (4) |
| S24L | 320 | 2020-03-08 | USA | 2020-05-13 | Australia: Victoria | USA (265) Australia (55) |
| D34E | 4 | 2020-04-28 | Australia: Victoria | 2020-04-30 | Australia: Victoria | Australia (4) |
| P36S | 5 | 2020-03-14 | USA: MA | 2020-04-04 | Australia: Victoria | USA (4) Australia (1) |
| P36L | 2 | 2020-03-22 | Australia: Victoria | 2020-04 | USA: Virginia | USA (1) Australia (1) |
| A51S | 2 | 2020-03-27 | USA: AK | 2020-03-31 | USA | USA (2) |
| V62L | 150 | 2020-01-22 | Hong Kong | 2020-05-11 | USA | USA (108) Australia (37) Hong Kong (3) Taiwan (1) India (1) |
| A65S | 13 | 2020-03-09 | USA: CA | 2020-04 | USA: Virginia | USA (10) Australia (2) Greece (1) |
| G66C | 2 | 2020-03-31 | USA: CT | 2020-06-01 | Bangladesh: Dhaka | USA (1) Bangladesh (1) |
| S67F | 4 | 2020-03-27 | USA: ID | 2020-05-10 | Australia: Victoria | USA (3) Australia (1) |
| S69L | 7 | 2020-03-10 | USA | 2020-04-06 | USA: Metairie, LA | USA (7) |
| L84S | 1000 | 2019-12-30 | China: Wuhan | 2020-05-27 | Australia: Victoria | USA (754) Australia (151) China (36) Thailand (13) Spain (13) India (9) Hong Kong (6) Greece (4) Puerto Rico (3) Tunisia (2) Taiwan (2) Uruguay (1) Kazakhstan (1) Japan (1) Germany (1) Egypt (1) Colombia (1) Bangladesh (1) |
| E110G | 2 | 2020-03-31 | USA | 2020-04-05 | USA | USA (2) |
| H112Q | 2 | 2020-03-11 | USA: WA | 2020-03-14 | USA: WA | USA (2) |
| V114F | 2 | 2020-03-22 | Australia: Victoria | 2020-03-24 | Australia: Victoria | Australia (2) |
| I121S | 6 | 2020-02-25 | China: Guangzhou | 2020-04-21 | USA: FL | USA (3) China (3) |

**Fig 8.**
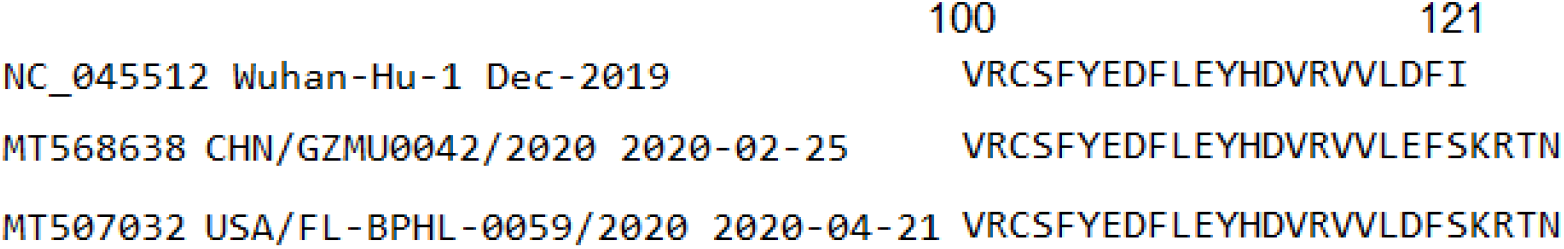
Alignment of protein ORF8 sequences having insertions with the GenBank accession numbers, isolate names and collected dates presented on the left. There is a high chance that the isolate CHN/GZMU0042/2020 (MT568638, collected in China) has transmitted to USA/FL-BPHL-0059/2020 (MT507032 in USA).

### Gene ORF10

The ORF10 protein has 38 AAs in length, appearing in 5,891 isolates with only 9 unique AA sequences. Among them, 5,872 sequences have no mutation or only synonymous mutations and the rest 19 sequences have nonsynonymous mutations. *No insertion and deletion mutations* are found in gene ORF10. Similar to ORF7b, this is a stable gene. There are 8 distinct nonsynonymous mutations, including I4L, A8V, S23F, R24L, R24C, A28V, D31Y and V33I. Those occurring in 2 sequences or more are presented in Table 9.

**Table 9.** Gene ORF10 - Nonsynonymous mutations occurring in 2 or more sequences

| **Mu** | **Total** | **First Date** | **First Country** | **Latest Date** | **Latest Country** | **Distribution** |
| --- | --- | --- | --- | --- | --- | --- |
| I4L | 7 | 2020-03-13 | USA: GA | 2020-04-15 | USA: Illinois | USA (7) |
| S23F | 5 | 2020-03-06 | Australia: Victoria | 2020-05-30 | Spain: Asturias | USA (2) Australia (2) Spain (1) |
| R24L | 2 | 2020-03-08 | USA: Massachusetts | 2020-03-09 | USA: WA | USA (2) |

## DETAILS OF MUTATIONS IN STRUCTURAL GENES: S, E, M AND N

### Gene S

The spike protein S has 1,273 AAs. The number of GenBank records having complete CDS of protein S is 4,434, with 259 unique AA sequences. Among them, 1,156 sequences have no mutation or synonymous mutations while other 3,278 sequences have deletion or nonsynonymous mutations. There are *no insertion mutation* among 4,434 sequences of protein S. There are 34 deletion mutations occurring in six sequences with *28 unique deletions*. Sequences in records MT012098 (India: Kerala State on 2020-01-27) and MT412290 (USA: WA on 2020-04-01) both have one deletion Y145-. Sequence in MT621560 (Hong Kong in 2020-03) has ten deletions continuously, consisting of N679-, S680-, P681-, R682-, R683-, A684-, R685-, S686-, V687- and A688-. Sequence in MT479224 (Taiwan on 2020-03-18) has 14 deletions, distributed in two deletion segments. The first segment includes nine deletions I68-, H69-, V70-, S71-, G72-, T73-, N74-, G75- and T76-while the second one includes five deletions Q675-, T676-, Q677-, T678-and N679-. Sequences MT474127 and MT460124 (both collected in USA: CA on 2020-03-27 and 2020-04-28, respectively) similarly have four deletions L141-, G142-, V143- and Y144-. The virus transmission may have happened between these two isolates but this needs further investigation. Alignment of these sequences is shown in Fig. 9.

**Fig 9.**
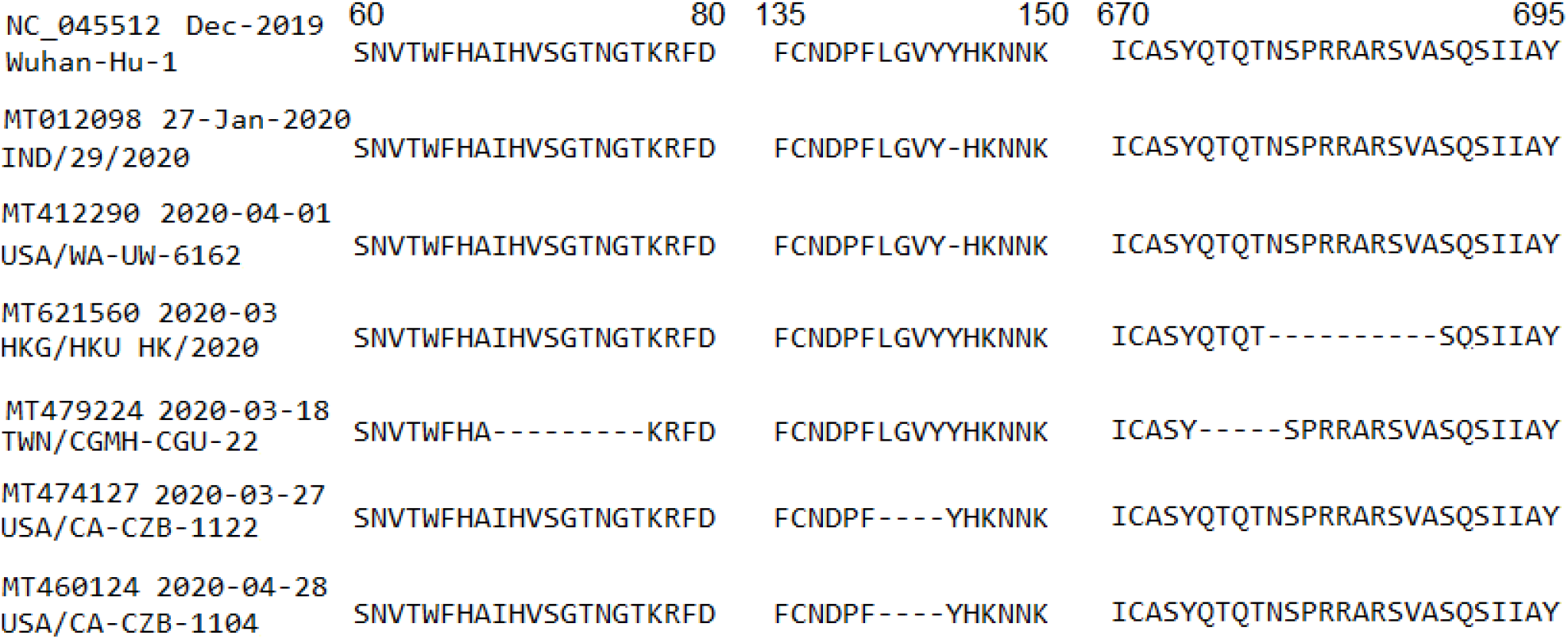
Deletions in protein S, which are all outside the RBD region (319-541), suggesting that the RBD may have been evolutionarily optimized for the purpose of binding to a host cell. The numbers on top show the residue positions in the protein. The GenBank accession numbers, collected dates and isolate names are presented on the left.

The number of nonsynonymous mutations in gene S is 3,711, with 240 distinct mutations. Mutations that occur in 10 or more cases are reported in Table 10. The number of synonymous mutations is 670, making a ratio between nonsynonymous versus synonymous mutations at 5.54. Among the nonsynonymous mutations, *mutation D614G* is extremely common as it happens in 3,089 sequences, majorly collected in USA (2340), India (210) and Australia (132). The first collected date of the D614G mutation cannot be identified precisely because some sequences deposited to the NCBI GenBank did not record the full date details. The current data show that either of the following sequences, which have the D614G mutation, was first collected: MT326173 in USA in 2020, or MT270104, MT270105, MT270108 and MT270109 all in Germany: Bavaria in 2020-01, or MT503006 in Thailand on 2020-01-04. It is however important to note that the first patient having the D614G mutation and his/her location may never be known because genome of that patient might not be sequenced and reported. Therefore, information reported here can support for further investigation.

**Table 10.** Gene S - Nonsynonymous mutations that occur in 10 or more sequences

| **Mu** | **Total** | **First Date** | **First Country** | **Latest Date** | **Latest Country** | **Distribution** |
| --- | --- | --- | --- | --- | --- | --- |
| L5F | 39 | 2020-03 | France | 2020-05-09 | Bangladesh: Narayanganj | USA (32) France (3) India (2) Italy (1) Bangladesh (1) |
| L54F | 12 | 2020-03-13 | USA: WA | 2020-05-24 | India: Ahmedabad | India (9) USA (2) Thailand (1) |
| D138H | 17 | 2020-04 | USA | 2020-06-01 | Bangladesh: Dhaka | USA (14) Australia (2) Bangladesh (1) |
| H146Y | 24 | 2020-03-20 | USA: UT | 2020-04-06 | USA | USA (24) |
| V483A | 11 | 2020-03-05 | USA: WA | 2020-04-05 | USA: WA | USA (11) |
| E554D | 14 | 2020-04 | USA | 2020-04 | USA | USA (14) |
| T572I | 13 | 2020-03 | France | 2020-06-03 | India: Ahmedabad | India (10) USA (2) France (1) |
| E583D | 13 | 2020-04 | USA: VA | 2020-06-03 | India: Ahmedabad | India (11) USA (1) Australia (1) |
| D614G | 3089 | 2020 / 2020-01 / 2020-01-04 | USA / Germany: Bavaria / Thailand | 2020-06-05 | India: Vadodara | USA (2340) India (210) Australia (132) Greece (77) France (61) Bangladesh (49) Germany (43) Japan (26) Poland (20) Czech Republic (20) Egypt (15) Taiwan (13) Thailand (11) Spain (10) Serbia (10) Italy (8) Puerto Rico (7) Netherlands (7) Tunisia (5) Turkey (4) Jamaica (3) China (3) Sri Lanka (2) Russia (2) Morocco (2) Kazakhstan (2) Hong Kong (2) South Africa (1) Peru (1) Nigeria (1) Israel (1) Colombia (1) |
| P681L | 16 | 2020-04 | USA: Virginia | 2020-04-03 | USA: CA | USA (16) |
| A829T | 37 | 2020-01-23 | Thailand | 2020-04-07 | Thailand | Thailand (37) |
| S939F | 11 | 2020-03-19 | USA: UT | 2020-04-15 | USA | USA (11) |
| P1263L | 10 | 2020-03 | France | 2020-04-09 | USA | USA (6) Greece (3) France (1) |

On the other hand, there are 37 A829T mutations that all occur in Thailand. The first case of this mutation was collected on 2020-01-23 and its latest case was on 2020-04-07. This may indicate that the first case had probably transmitted to other cases having the same mutation A829T in Thailand. Alternatively, mutations H146Y (24 cases), V483A (11 cases), E554D (14 cases), P681L (16 cases) and S939F (11 cases) all occur only in USA or mutation L8V (4 cases) occurs only in Hong Kong (refer to the spreadsheet data, which can be found in the Data Availability section).

We identify the RBD region within the residue range Arg319-Phe541 of protein S based on a study in^38^. In the RBD region only, the number of nonsynonymous mutations is 53 and that of synonymous is 46, making a ratio of 1.15. This is much smaller than the ratio of 5.54 for the entire gene S, suggesting that the RBD region may have been optimized for binding to a receptor of a host cell. This is complemented by Fig. 9 showing all deletion mutations in gene S being outside the RBD region. Note that the difference of these ratios is partly due to the large number of D614G mutations (3,089), which is outside the RBD region.

Table 11 summarizes nonsynonymous mutations in the RBD region occurring in 2 or more sequences. Notable mutation in this region is V483A occurring in 11 isolates all collected in USA. The first and latest collected dates of these isolates were respectively 2020-03-05 and 2020-04-05, suggesting that the first isolate may have spread to others having the same mutation V483A. Likewise, the mutation G476S occurs in 6 isolates all collected in USA: WA from 2020-03-10 to 2020-03-25. Alternatively, the mutation Y453F occurs in 5 sequences all in Netherlands but the first collected date was on 2020-04-25 and the latest collected date was on 2020-04-29. These dates are too close, indicating that all the reported Y453F cases may have been infected from another case, whose genome had not been sequenced and reported to the NCBI GenBank. It is important to note that all the transmission implications need further investigation with more data from other aspects such as travel history, physical contacts and so on.

**Table 11.** Gene S - RBG region only - Nonsynonymous mutations that occur in 2 or more sequences

| **Mu** | **Total** | **First Date** | **First Country** | **Latest Date** | **Latest Country** | **Distribution** |
| --- | --- | --- | --- | --- | --- | --- |
| V367F | 5 | 2020-01-22 | Hong Kong | 2020-05-06 | Netherlands | Hong Kong (3) USA (1) Netherlands (1) |
| R408I | 2 | 2020-01-27 | India: Kerala | 2020-05-02 | Egypt | India (1) Egypt (1) |
| Y453F | 5 | 2020-04-25 | Netherlands | 2020-04-29 | Netherlands | Netherlands (5) |
| G476S | 6 | 2020-03-10 | USA: WA | 2020-03-25 | USA: WA | USA (6) |
| V483A | 11 | 2020-03-05 | USA: WA | 2020-04-05 | USA: WA | USA (11) |
| G485R | 2 | 2020-02-06 | China | 2020-02-06 | China | China (2) |
| S494P | 3 | 2020-03-20 | USA: Michigan | 2020-03-20 | USA: Michigan | USA (3) |
| H519Q | 2 | 2020-03-15 | USA: WA | 2020-04-27 | Australia: Victoria | USA (1) Australia (1) |
| A520S | 3 | 2020-03-13 | USA: WA | 2020-04-26 | USA: AK | USA (3) |

In gene S, 164 nonsynonymous mutations (69 unique) are likely to make changes in protein secondary structure. In the RBD region, S477G [CBCTT(S)CCCCC -> CBCTT(S)CCCEC], P479L [CTTSC(C)CCCCC -> CTTSC(C)CECCC], V483A [CCCCC(C)CTTTC -> CCCCC(C)CCTTC], and F486L [CCCCT(T)TCBCS -> CCCEC(T)TCBCS] are four mutations having protein structure change potential.

On the other hand, 184 nonsynonymous mutations (58 unique) have the relative solvent accessibility change potential. Of these, only three mutations are in the RBD region: V483A [eebbb(b)bebeb-> eebbb(b)bbbeb], G485R [bbbbb(e)bebee -> bbbbb(b)bebee], and F486L [bbbbe(b)ebeeb -> bbbbb(e)ebeeb]. Two mutations *V483A and F486L* are thus likely to make changes in both protein secondary structure and relative solvent accessibility in the RBD region.

For the entire protein S, 134 nonsynonymous mutations (48 unique) have both structure and solvent accessibility change potentials. These mutations occurring in 2 or more sequences are reported in Table 12. *Mutation H146Y* occurs in 24 cases and *mutation P681L* occurs in 16 cases, which are all collected in USA. The most common mutation *D614G does not have the potential* to change either protein secondary structure or relative solvent accessibility.

**Table 12.**
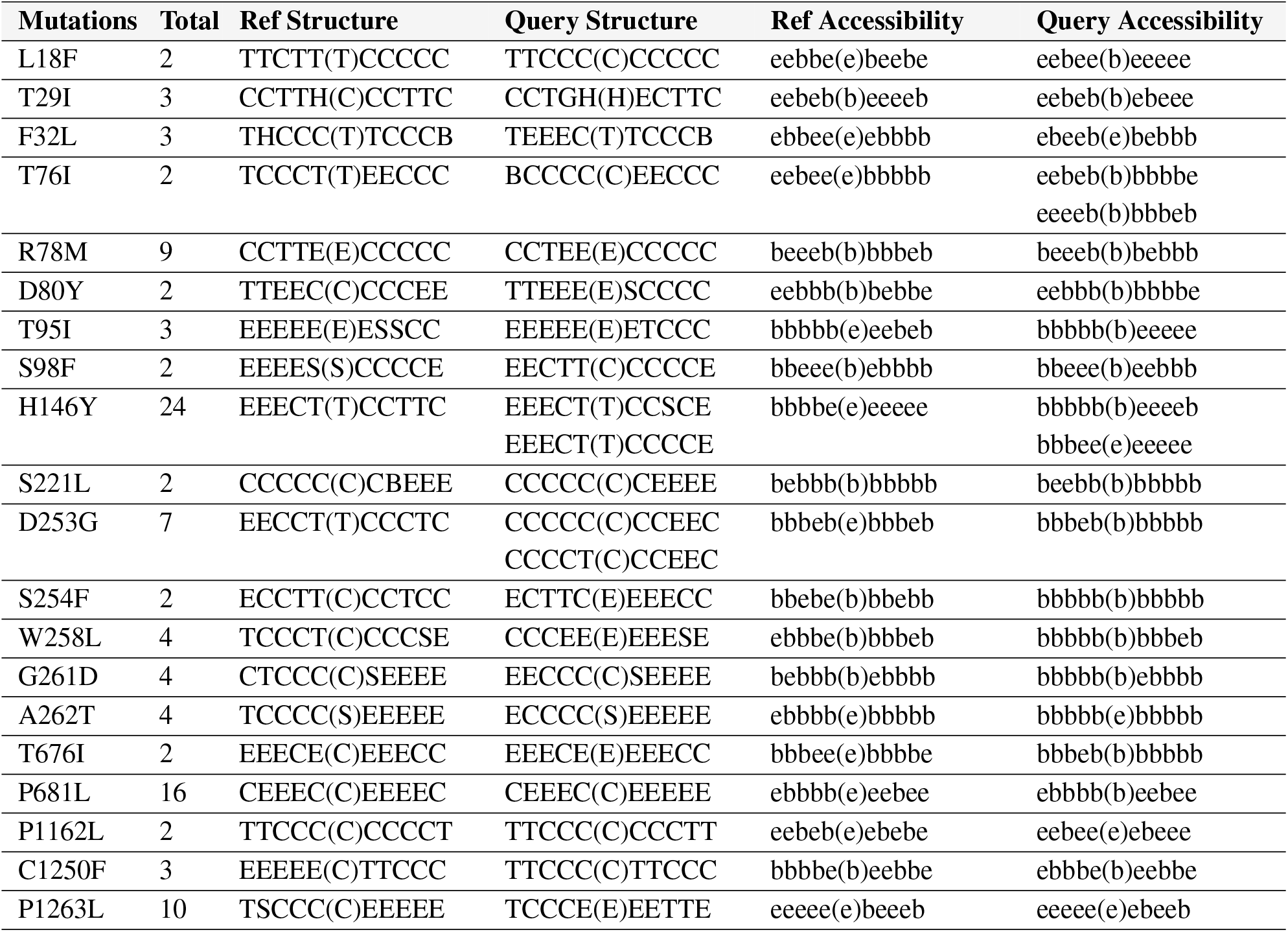
Gene S - Nonsynonymous mutations that have both structure and solvent accessibility change potentials occuring in 2 or more sequences. The “Query Structure” (and “Query Accessibility”) shows the unique structure (and accessibility) changes based on on prediction results. Structure letter in parentheses is the predicted structure of the residue at the corresponding mutation position. Five letters before and after parentheses are structures of neighbouring residues. Likewise, letter “b” or “e” in parentheses shows the accessibility status of the residue at the mutation position.

### Gene E

The envelope protein E has 75 AAs, found in 5,852 GenBank records with 15 unique AA sequences. Among them, 5,824 sequences have no mutation or only synonymous mutations while 28 sequences have nonsynonymous mutations. Gene E is thus relatively stable and could be targeted for vaccine and drug development. This is supported by the fact that *no insertion or deletion mutations* are found within gene E. There are *14 distinct nonsynonymous mutations* in gene E and those occur in 2 or more sequences are presented in Table 13.

**Table 13.** Gene E - Nonsynonymous mutations occurring in 2 or more sequences

| Mu | Total | First Date | First Country | Latest Date | Latest Country | Distribution |
| --- | --- | --- | --- | --- | --- | --- |
| F26L | 2 | 2020-04-06 | USA | 2020-04-06 | USA: New Orleans, LA | USA (2) |
| S68F | 5 | 2020-03-03 | Australia: Victoria | 2020-05-22 | Kenya | USA (2) Australia (2) Kenya (1) |
| P71L | 7 | 2020-03-19 | USA: WA | 2020-04-30 | USA: CA | USA (7) |
| L73F | 4 | 2020-03-22 | Australia: Victoria | 2020-04-07 | Australia: Victoria | Australia (3) USA (1) |

Five distinct nonsynonymous mutations in gene E have protein structure change potential: S68C, S68F, P71L, D72Y and L73F. Alternatively, 4 distinct mutations have potential to change relative solvent accessibility: L37H, L37R, D72Y and L73F. Therefore, *D72Y and L73F* are two mutations in gene E that have a potential to change both protein structure and solvent accessibility.

### Gene M

The M protein has 222 AAs and its complete CDS appears in 5,677 GenBank records, with 37 unique AA sequences. There are 5,557 sequences having no mutation or only synonymous mutations while other 120 sequences have nonsynonymous mutations. *No insertion or deletion mutations* are found in gene M. The number of distinct nonsynonymous mutations in gene M is 37, with those occurring in 5 or more sequences shown in Table 14. Among these, 10 mutations are likely to make changes in protein secondary structure: C64F, A69S, A69V, V70F, N113B, R158L, V170I, D190N, D209Y and S214I. Alternatively, 6 mutations have the solvent accessibility change potential: N113B, P123L, P132S, H155Y, D190N and T208I. *N113B and D190N* are thus two mutations having potential to change both protein structure and solvent accessibility in gene M.

**Table 14.** Gene M - Nonsynonymous mutations occurring in 5 or more sequences

| Mu | Total | First Date | First Country | Latest Date | Latest Country | Distribution |
| --- | --- | --- | --- | --- | --- | --- |
| A2S | 5 | 2020-01-29 | China: Beijing | 2020-05-11 | USA | USA (4) China (1) |
| D3G | 20 | 2020-03-05 | USA: SC | 2020-03-27 | Australia: Victoria | Australia (13) USA (4) Thailand (1) Kazakhstan (1) Italy (1) |
| V70F | 13 | 2020-03-14 | USA | 2020-04-01 | Australia: Victoria | USA (12) Australia (1) |
| V70I | 9 | 2020-03-10 | USA: WA | 2020-04-06 | USA: WA | USA (9) |
| R146H | 8 | 2020-04-09 | Australia: Victoria | 2020-04-15 | Australia: Victoria | Australia (8) |
| T175M | 18 | 2020-03 | France | 2020-04-01 | Hong Kong | USA (5) Australia (5) Czech Republic (4) Russia (1) Hong Kong (1) Germany (1) France (1) |

### Gene N

The N protein has 419 AAs and its complete CDS appears in 5,281 isolates, with 178 unique AA sequences. Among them, 4,315 sequences have no mutation or only synonymous mutations while the rest 966 sequences have deletions or nonsynonymous mutations. There are *no insertion* in gene N. The sequence in MT434815 (collected in USA: NY on 2020-03-09) has three sequential deletions at Q390-, T391- and V392-while the sequence in MT370992 (USA: NY on 2020-03-20) has six sequential deletions at T366-, E367-, P368-, K369-, K370- and D371-. Two other sequences MT605818 and MT560525 (both collected in Turkey on 2020-04-16) have three sequential deletions at R195-, N196- and S197-. There are 1,927 nonsynonymous mutations with 156 distinct ones and those occurring in 10 or more sequences are presented in Table 15. Notable mutations are R203K occurring in 871 sequences and G204R occurring in 433 sequences. There are 15 mutations in this protein having the potential to change both protein structure and solvent accessibility, including G18V, D22Y, G34W, R40C, R40L, R185C, A211S, P365H, T391I, T393I, A398S, D399E, D399H, D401Y and D402Y.

**Table 15.** Gene N - Nonsynonymous mutations occurring in 10 or more sequences

| **Mu** | **Total** | **First Date** | **First Country** | **Latest Date** | **Latest Country** | **Distribution** |
| --- | --- | --- | --- | --- | --- | --- |
| P13L | 93 | 2020-02-27 | South Korea | 2020-05-14 | Australia: Victoria | Australia (43) India (23) USA (10) Timor-Leste (3) South Korea (3) Malaysia (3) Guam (3) Taiwan (2) Thailand (1) Jamaica (1) France (1) |
| G18C | 10 | 2020-04-22 | USA: Michigan | 2020-04-29 | USA: Michigan | USA (10) |
| D22G | 10 | 2020-03-18 | Australia: Victoria | 2020-03-30 | Australia: Victoria | Australia (10) |
| S183Y | 15 | 2020-03-17 | USA: NY | 2020-04-08 | USA: Massachusetts | USA (14) Australia (1) |
| S194L | 102 | 2020-01-29 | USA: MA | 2020-06-05 | India: Gandhinagar | India (79) USA (10) Australia (6) Bangladesh (3) Hong Kong (2) Taiwan (1) Germany (1) |
| S197L | 44 | 2020-02-26 | Spain: Valencia | 2020-04-16 | Australia: Victoria | Australia (26) Spain (10) USA (5) Greece (2) Kazakhstan (1) |
| S202N | 25 | 2020-01-30 | China: HuaShang | 2020-05-27 | Australia: Victoria | USA (10) India (8) Tunisia (2) China (2) Australia (2) Bangladesh (1) |
| R203K | 871 | 2020-02 | Germany | 2020-06-01 | Bangladesh: Dhaka | USA (256) Australia (203) Greece (122) Bangladesh (88) Japan (38) Czech Republic (34) Poland (26) Germany (18) India (12) Taiwan (10) Turkey (8) France (8) Thailand (6) Serbia (6) Italy (6) Hong Kong (6) Spain (4) Russia (4) Sri Lanka (2) Puerto Rico (2) Peru (2) Nigeria (2) Morocco (2) Kazakhstan (2) Israel (2) China (2) |
| G204R | 433 | 2020-02 | Germany | 2020-06-01 | Bangladesh: Dhaka | USA (126) Australia (101) Greece (61) Bangladesh (44) Japan (19) Czech Republic (17) Poland (13) Germany (9) India (6) Taiwan (5) Turkey (4) France (4) Thailand (3) Serbia (3) Italy (3) Hong Kong (3) Spain (2) Russia (2) Sri Lanka (1) Puerto Rico (1) Peru (1) Nigeria (1) Morocco (1) Kazakhstan (1) Israel (1) China (1) |
| T205I | 18 | 2020-01-29 | China: Beijing | 2020-04-05 | Australia: Victoria | USA (9) Greece (5) Australia (3) China (1) |
| A208G | 26 | 2020-03 | USA: VA | 2020-04-09 | USA: FL | USA (24) Taiwan (1) Australia (1) |
| T362I | 17 | 2020-04 | USA: Virginia | 2020-04-29 | India: Ahmedabad | USA (16) India (1) |

## DISCUSSIONS

The proposed analysis approach has been able to detect all point mutations so far of SARS-CoV-2 and report them in a spreadsheet dataset, which can be found in the Data Availability section. The generated data can facilitate investigations about the virus in many perspectives. For example, using the mutations found, we can observe the possible virus transmissions between patients. This is showcased through Figs. 5, 6 and 8 where similar mutations are detected from different isolates. In Fig. 5 for instance, a group of isolates have similar deletion mutations at positions M85-, V86- or K141-, S142-, F143-in protein ORF1ab. Genome sequence of isolate USA-CA6/2020 collected in USA on 2020-01-27 is found to be the first having these consecutive mutations. There could be a connection between this isolate with other isolates obtained in USA, Greece and Australia. Information presented in Tables 2-15 is extracted from the generated mutation data. In these summary tables, “First Date” and “First Country” information is useful for identifying when and where the mutations were possibly originated while the “Distribution” information shows how such mutations have spread to different countries.

The generated mutation data also allow us to observe the evolution of the virus. We can point out which mutations are still active or no longer active based on the “Latest Date” information presented in Tables 2-15. For example, in gene S (Table 10), the latest date of D614G was on 2020-06-05 (same as the latest collection date of the entire genome dataset used in this study), indicating that this mutation is still active. The latest date of P681L was on 2020-04-03, showing that this mutation may no longer occur. Mutations are still active mean they have evolutionarily adapted to varying environments while inactive mutations may have not. This kind of information is useful for further research on vaccine and drug development as ongoing changes of the viral proteins need to be focused and addressed rather than inactive mutations. Our analysis shows that the G form at location 614 in protein S becomes increasingly popular compared to the D form. Among 4,434 sequences of the S protein, 3,089 sequences have the mutation D614G, taking 69.67%. This number has considerably increased compared to 31.5% in the previous analysis in^22^ on a dataset downloaded on 2020-03-22.

Through the mutation analysis, we are able to detect regions of the viral genomes that are stable and can be targeted for vaccine and drug development, such as those coding for proteins E, M, ORF6, ORF7a, ORF7b and ORF10. In addition, this study presents results obtained by the use of deep learning recurrent neural networks for protein secondary structure and solvent accessibility predictions. These results are useful for further research on SARS-CoV-2 protein structure changes. In particular, among 3,089 D614G mutations, our prediction results show that none of these mutations is likely to make changes in the protein secondary structure and relative solvent accessibility. This mutation has attracted much attention of researchers as it may affect the virulence and infectivity of the virus and our finding has contributed to understanding this mutation.

## CONCLUSIONS

Analysing the virus genome sequences and their proteins is crucial for understanding the virus and proposing appropriate approaches to respond to and control the pandemic. This paper has reported all point mutations of SARS-CoV-2 since the virus’ s first genomes were obtained in December 2019. A SARS-CoV-2 mutation spreadsheet dataset is built using a large number of genome sequences (6,324) obtained across 45 countries. This dataset can enable scientists to monitor the evolution and spread of the virus although the use of these data needs to be corroborated with patients’ clinical data and travel history for substantiated confirmations. We also predict the secondary structure and relative solvent accessibility of the virus proteins to evaluate whether the detected mutations have a potential to change the virus characteristics. The mutation D614G in protein S is unlikely to change either protein secondary structure or relative solvent accessibility based on the prediction results. These protein secondary structure and solvent accessibility change potentials are predicted results based on deep learning recurrent neural networks, which need to be experimentally verified. They however provide important insights about the virus and prompt further experimental biochemistry and molecular biology research into the genomic regions of these mutations. A future work focusing on impacts of the mutations on protein functions would be worth investigating. The function impacts can be determined via wet lab experiments or using low-cost computational tool such as PROVEAN^39^.

## DATA AVAILABILITY

The dataset generated and analysed during the current study is available in the bioRxiv repository, https://www.biorxiv.org/content/10.1101/2020.07.10.171769v2.supplementary-material.

